# Modeling Spatial Correlation of Transcripts With Application to Developing Pancreas

**DOI:** 10.1101/391433

**Authors:** Ruishan Liu, Marco Mignardi, Robert Jones, Martin Enge, Seung K Kim, Stephen R Quake, James Zou

## Abstract

Recently high-throughput image-based transcriptomic methods were developed and enabled researchers to spatially resolve gene expression variation at the molecular level for the first time. In this work, we develop a general analysis tool to quantitatively study the spatial correlations of gene expression in fixed tissue sections. As an illustration, we analyze the spatial distribution of single mRNA molecules measured by in situ sequencing on human fetal pancreas at three developmental time points 80, 87 and 117 days post-fertilization. We develop a density profile-based method to capture the spatial relationship between gene expression and other morphological features of the tissue sample such as position of nuclei and endocrine cells of the pancreas. In addition, we build a statistical model to characterize correlations in the spatial distribution of the expression level among different genes. This model enables us to infer the inhibitory and clustering effects throughout different time points. Our analysis framework is applicable to a wide variety of spatially-resolved transcriptomic data to derive biological insights.

## Introduction

The spatial heterogeneity of gene expression has attracted much attention in disease, medicine and developmental studies. Understanding transcriptional heterogeneity provides critical information to interpret biological processes and to develop clinical therapies^1,2^. For decades, immunohistochemistry has been the workhorse for studying the protein expression in tissue samples. Although robust, this method is limited to the study of few proteins at time and it is sometimes hampered by the poor performance of the antibody used. On the contrary, transcriptome measurements are now performed genome-wide either with bulk measurements of the tissue of interest or by analysis of single cells extracted from the tissue. The spatial resolution is lost in both approaches^3,4^.

Recently, in situ sequencing and other fluorescent in situ hybridization (FISH) - based methods were developed, which enabled high-resolution spatially resolved transcriptomic studies^5–9^. These technologies image and detect RNA molecules directly in tissue samples, thus maintaining the spatial information with high resolution. In contrast to the rapid growth of in situ transcriptomic technologies, the computational analysis on the spatial transcriptomic data is still in its infancy. Most studies are carried out in a non-quantitative manner or only provide preliminary statistics^10,11^. Recent methods such as Spatia1IDE aim to identify individual genes that are spatially varying but do not model gene-gene spatial correlations^12^. Many important spatial characteristics remain unexplored, and the poor quantification becomes a severe problem, especially when comparison across different time points is required like in developmental studies. Therefore computational methods to explore these novel datasets are needed.

We develop a general analysis tool to explore and quantitatively study the spatial distribution of gene expression data generated by in situ transcriptomic methods. We demonstrate our approach by exploring spatial transcriptomic data generated by in situ RNA sequencing of human fetal pancreas tissues of different ages 80, 87 and 117 days post-fertilization. A density profile-based method is developed to capture the relation between gene expression and other biological targets such as cell nuclei and forming pancreatic islets of the Langerhans. A statistical model is built to characterize the spatial interactions among the expression of different genes. This new tool allows us to model and measure inhibition or clustering effects between transcripts expressed by different cells in the tissues. As a broadly new perspective in development studies, we show that our method can be used as an exploratory tool to identify spatial gene interactions of potential importance in the development of the pancreas. Our tool is publicly available at https://github.com/RuishanLiu/Gene-Spatial.

## In Situ RNA Sequencing

In situ techniques enable us to spatially resolve gene expression by performing molecular reactions directly in fixed cells and tissue sections ^13^. The techniques achieve high multiplexing by two main strategies: combinatorial decoding or sequencing-based readout. Combinatorial decoding methods, typically exploited by fluorescent in situ hybridization (FISH) assays, use fluorescently labeled probes in multiple combinations to distinguish a large number of different targets, each one corresponding to a specific color combination in a predetermined color-coding scheme^14,15^. Sequencing-based readouts for in situ assays build on biochemical methods developed for parallel DNA sequencing in next-generation sequencing (NGS) platforms and apply them to a molecular substrate that is generated directly in fixed tissue^5,6^.

The gene expression data analyzed in this work are generated by a combination of these methods. Specifically, single RNA molecules are amplified as previously described by Ke et al. (In situ sequencing)^5^ using a gene-tiling approach. Each gene is targeted by 1 to 13 different cDNA primers which hybridize at different positions along the length of the mRNA. This increases the probability to successfully reverse transcribe the gene. Each primer is coupled with a gene-specific barcoded padlock probe which is ligated to the cDNA and subsequently amplified via rolling-circle amplification (RCA). The molecular barcodes associated with each transcript are then decoded by sequential hybridization of fluorescence probes following a combinatorial decoding scheme. The protocol used to stain the tissues is detailed in the Supplementary Note 1 along with the list of targeted genes and the probe sequences.

Every round of hybridization is carried out using four oligonucleotide probes, each one labeled with a distinct fluorophore, which are hybridized to the amplified cDNA molecules directly on a section of pancreatic tissue. The total barcoding space results in 4^3^ = 64 possible combinations. Here we assign 25 combinations to transcripts from 25 different genes, and leave the remaining 39 combinations as negative controls to assess sequencing quality. The targeted genes list comprises a number of marker genes for endocrine cells (alpha, beta and delta cells), transcription factors implicated in differentiation of the endocrine cells and genes expressed in mesenchymal cells at different levels during pancreas development. The data are collected in samples from three developmental ages 80, 87 and 117 days post-fertilization.

The RNA molecules of an entire tissue section undergo three rounds of staining and imaging as diffraction-limited spots in its native cellular context together with a nuclear staining (DAPI). The collected images are processed as described previously^5,16^ and a detailed description of the image processing can be found in Supplementary Note 1. The intensity values are extracted from each individual diffraction-limited signal, where the fluorescent probes appear as bright round spots. The raw data quality is potentially affected by the influence of neighboring fluorescence, misalignment between the three rounds of imaging and camera noise. For example, 7% of detected RNA molecules (11,611 out of 159,716) are labeled by the 39 negative control combinations at day 87. To carry out quality control, we define a quality metric as the averaged confidence of fluorescence and filter out all the detected transcripts with a quality lower than 55%. After the processing, 32% of the data are discarded and as low as 2% of the rest (2,647 out of 108,430) have meaningless labels. This is in accordance with the accuracy of the method as previously described ^5^.

For every image, the position (x, y coordinates) of each segmented nuclei and detected transcript as well as the transcript identity are recorded and can be plotted like in Figure 1a where the spatial distribution of three mRNA transcripts somatostatin (SST), glucagone (GLUC) and insulin (INS) is shown in 2D coordinates (x, y). At day 117, for example, 159,716 RNA molecules for the 25 types of genes are detected. The slice of pancreas has 50,147 cells in total, and the nucleis positions are illustrated in Figure 1b. We first focus our computational analysis on data from day 117, since that represents the highest quality data. Then in the *Temporal Analysis* Section, we discuss how we integrate data from earlier time points to model temporal differences in spatial expression.

**Figure 1:**
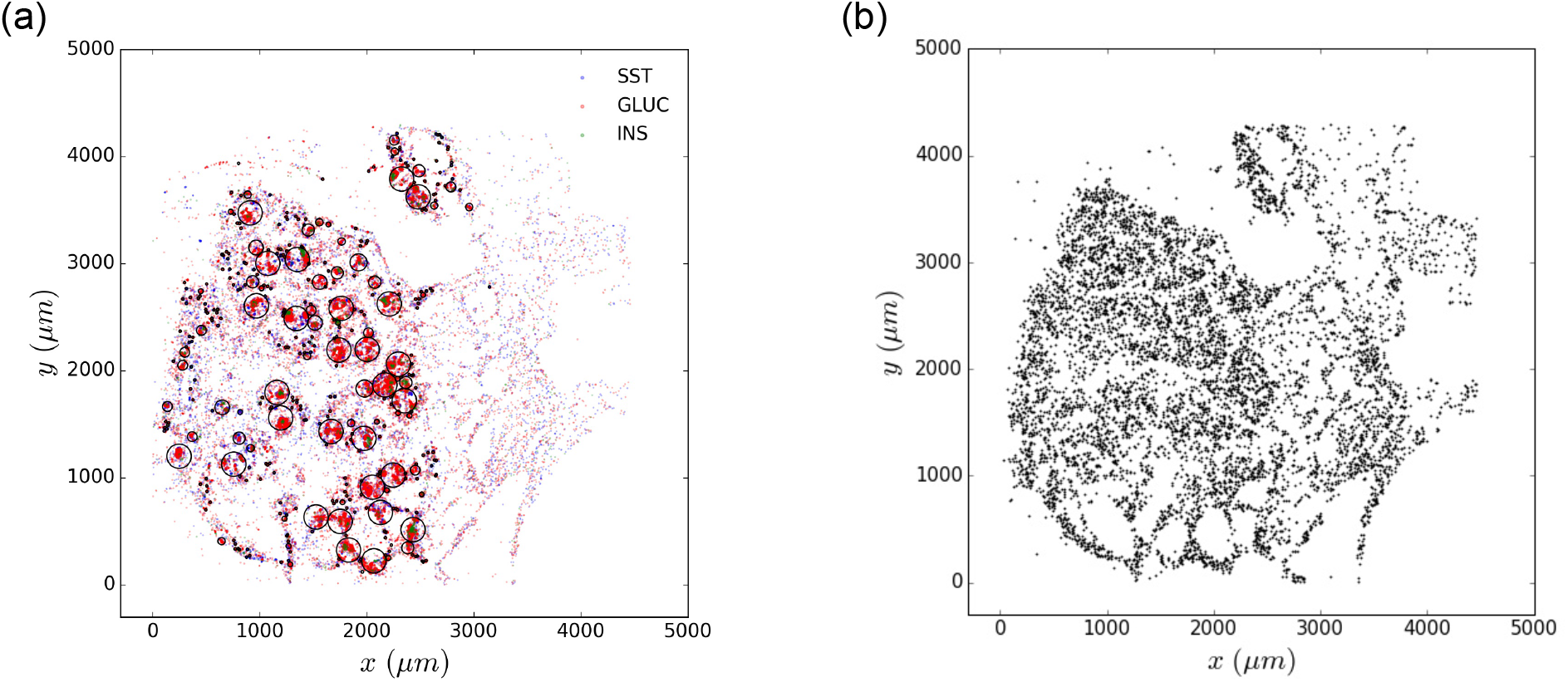
In situ sequencing. The sample is from fetal pancreas at age 117 days post fertilization. (a) Detected SST, GLUC and INS transcripts are plotted on xy coordinates. Computationally identified pancreatic islets are identified by black circles. (b) Identified and segmented nuclei are plotted on xy coordinates.

## Computational Analysis

### Identification of Pancreatic Islets

The pancreas is composed of a hormone-producing compartment (endocrine pancreas) and a digestive enzyme-producing one (exocrine pancreas). In the developed organ, the endocrine compartment is organized in discrete units, known as islets of Langerhans. These are clusters of hormone- producing cells mostly alpha, beta and delta cells which produce respectively Glucagone (GLUC), Insulin (INS) and Somatostatin (SST). Endocrine and exocrine areas have different physiological functions and cell type composition, thus the study of spatial properties requires identification of the morphological context.

Here, endocrine islets in the process of formation are identified using a clustering algorithm that we developed (Algorithm 1 in Supplementary Note 3). SST, GLUC and INS transcripts are used as marker genes for identification of endocrine cells and pancreatic islets. For the convenience of computation thereafter, all the endocrine islets are assumed to be circular. Real boundaries for islets could be approximated by circles. As shown in Figure 1a, the algorithm is able to identify large islets as well as single cell exocrine regions. The distribution of identified islets size is provided in Supplementary Figure 2. The wide variation in islets diameter has been reported in early as well as more recent studies. However, in the fetal samples the maximum diameter size of islets is smaller than previously reported in the adult pancreas (300 μm)^17,18^.

**Figure 2:**
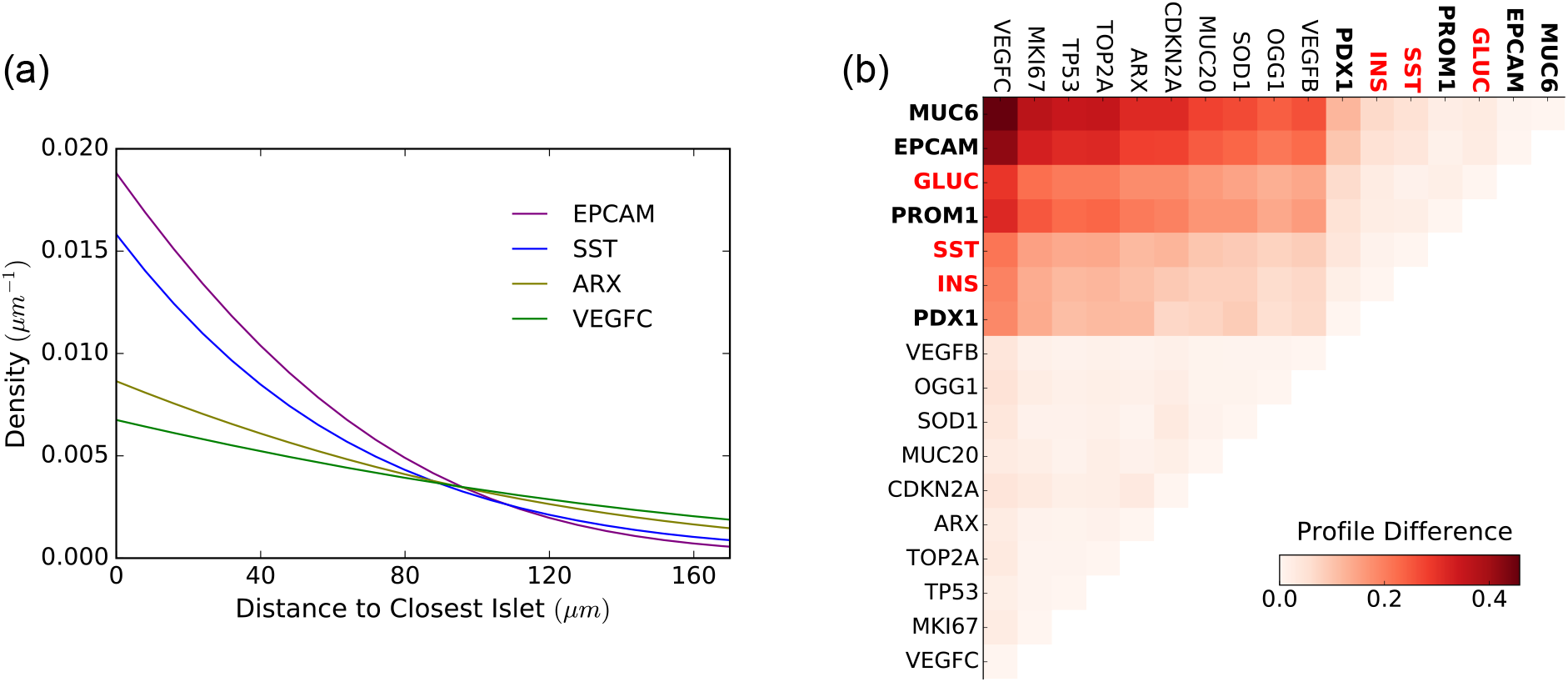
Islets-related density profile. Here the sample is collected at age 117 days post fertilization. (a) An example of two different density profiles for four selected genes in respect to pancreatic islets. EPCAM and SST show a higher density closer to islets compared to ARX and VEGFC. (b) The difference between the density profiles is calculated and plotted as heatmap. Two groups of genes can be identified. In bold are the genes belonging to group one. In red are the genes used to identify the islets and therefore expected to be found closer to them.

### Density Profile-based Analysis

To capture the relation between transcripts and other morphological features of the tissue such as nucleis position or developing pancreatic islets, we carried out a density profile-based analysis. The density profiles are calculated based on kernel density estimation with linear combination correction ^19^. The difference between two density profiles is characterized by KL divergence. See Supplementary Note 2 for more details.

First, we focused on the spatial relationship between endocrine islets and the transcripts which are outside the islets, in order to identify genes whose expression resulted enriched in proximity of the forming islets. These genes may be directly involved in the differentiation of endocrine cells or be constitutively expressed in the cells nearby. Only the most abundant genes which have at least 100 counts are examined (17 out of 25 genes). At day 117, these are genes which contribute at least 0.1% of the total reads. As an example, the density profile of some transcripts with respect to their distance to the closest islets boundary on day 117 is illustrated in Figure 2a. For each gene pair, the KL divergence of density profiles indicates the difference between the spatial distributions of two genes outside endocrine islets and is plotted in Figure 2b. The larger the difference is, the more distinct the two density profiles are.

Based on the KL divergence of density profiles we identify two groups of genes with distinct density distribution profile from each other. The two groups are highlighted in Figure 2b, where MUC6, EPCAM, GLUC, PROM1, SST, INS and PDX1 form Group 1, marked in bold, and the rest of genes form Group 2. Within Group 1, INS, GLUC and SST are markers for endocrine cells and are expected to be found within or in proximity of pancreatic islets. PDX1 is a transcriptional activator of several genes, including insulin and somatostatin, and is involved in the early development of the pancreas, which plays a major role in glucose-dependent regulation of insulin gene expression ^20^. EPCAM is an antigen expressed in epithelial cells and in stem cells ^21^, and PROM1 (CD133) is a surface antigen found in progenitor and stem cells in the mouse and human pancreas ^22,23^. Their distribution profile clustered together with endocrine cell markers which may indicate a potential role for these genes in the differentiation of progenitor cells into endocrine cells. The MUC6 gene transcribes a glycoprotein belonging to the mucin family, a class of protein which are found in many epithelial tissues. Increasing expression of MUC6 has been observed during development of several human organs including pancreas, but its role has not been well defined yet ^24,25^.

Similarly our spatial analysis of gene expression could be carried out on other tissue features such as the nuclei. In this case, the density profile captures how likely it is to find a transcript as we move further away from the cell nucleus. As an example, the density profiles of some transcripts with respect to the closest nucleus at day 117 are plotted in Figure 3. However, because automatic nuclei and cell segmentation is particularly difficult in our data and in in situ methods in general, the retrieved nuclei locations may not be accurate.

**Figure 3:**
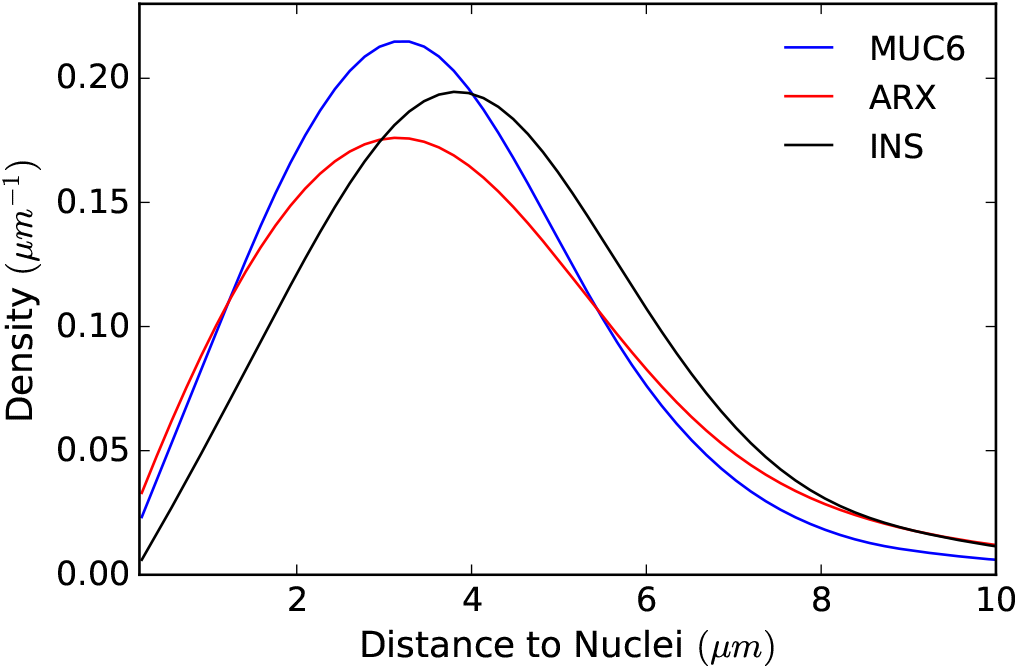
Nuclei-related density profile. An example of three density profiles for three genes in respect to their closer nuclei. Genes are assigned to the closer nucleus identified by segmentation of the DAPI staining images. Here the sample is collected at age 117 days post fertilization.

### Temporal Analysis

We then asked whether the transcriptional density profile observed for sample aged 117 days differs from profiles of samples at earlier time points. A difference could be indicative of transient gene expression in the vicinity of the endocrine cells and thus identify genes involved in development of specific cell types at specific time points.

We found that the distinction between the two groups of transcripts identified in sample 117 day shows a temporal trend, becoming larger at later time points — distinction is small at day 80 in Figure 4a, moderate at day 87 in Figure 4b, and most obvious at day 117 in Figure 2b. The density profiles of marker genes which demarcate forming islets cluster more and more during development and the identified groups of genes separate markedly from each other. Thus as the tissue structures (pancreatic islets) become more and more evident with time so does their gene expression profile distribution.

**Figure 4:**
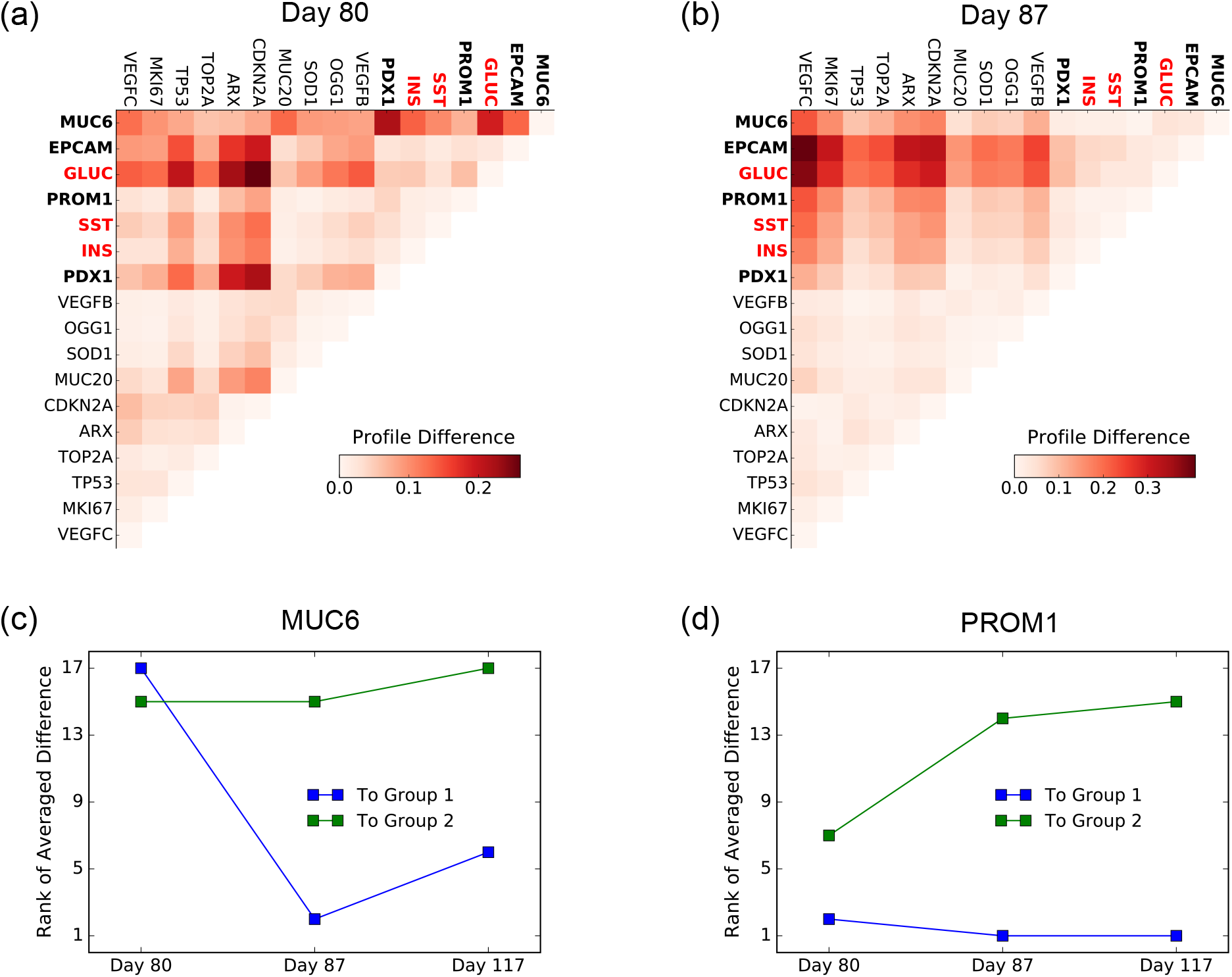
Islets-related temporal analysis of density profiles. (a-b) The difference between the density profiles for samples age 80 and 87 days after fertilization is calculated and plotted as heatmap. The two groups of genes identified in sample age 117 days are still evident but to a lesser extent. In bold are the genes belonging to group one. In red are the genes used to identify the islets and therefore expected to be found closer to them. The rank of average difference from the two groups can be plotted for each single gene. Here the difference at the three time points is shown for (c) MUC6 and for (d) PROM1.

In addition our analysis can reveal temporal changes in expression distribution for individual genes, which could also be of potential biological interest. To do so we rank the averaged difference between one gene and the genes in one of the two groups. For instance, MUC6 is found to become closer to Group 1 during development, as shown in Figure 4c. At day 80, MUC6 is farthest to Group 1 among 17 genes, but is the fifth closest to Group 1 at day 117. This suggests that MUC6 might play a particularly dynamic role in islet development. PROM1 is found to become more distinct to Group 2 across the time, as depicted in Figure 4d.

### Statistical Model for Spatial Correlations

To characterize the spatial distribution of the expression level among different genes, we carried out the analysis based on a statistical model. Within the range of imaged tissue, the spatial transcriptome is characterized by the likelihood ratio 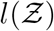, where 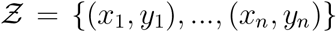 is the set of transcripts positions. Here we use the multitype Strauss process model^26,27^

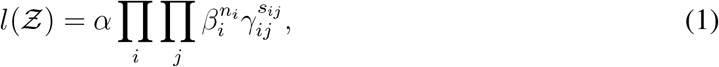

where *α* is a normalizing constant, *β_i_* indicates the intensity of type *i* transcripts, *γ_ij_* denotes the spatial correlations between type *i* and type *j* transcripts, *n_i_* is the number of type i transcripts in 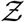 and *s_ij_* is the number of type *j* transcripts in the neighbor of type i within radius *r*. The correlations are fully described by *³_ij_*. The case 0 < *γ_ij_* < 1 indicates an inhibition effect between the expression of type *i* and type *j* genes, and *γ_ij_* > 1 represents a clustering effect. If *γ_ij_* = 1 for all *i* and *j*, Eq. (1) gives a Poisson process with intensity *β_i_* for type i transcripts. To increase the resolution of our analysis we applied this model within each cluster of endocrine cells to test for clustering or inhibition effects among these cells. To capture the short-range interactions, the radius is set to be 20 *μm*, twice as the averaged nuclei spacing likely describing interactions between neighboring cells or genes co-expressed in the same cell. Only genes detected at a threshold level of 500 transcripts within endocrine islets are analyzed and the results are summarized in Table 1.

**Table 1:** Number of transcripts inside endocrine islets for 7 genes at age 117 days post fertilization.

|  | **GLUC** | **SST** | **INS** | MUC6 | EPCAM | PROM1 | ARX |
| --- | --- | --- | --- | --- | --- | --- | --- |
| Total Number in Islets | 37171 | 11611 | 1904 | 4665 | 1056 | 774 | 697 |

Most spatial correlations *γ_ij_* among genes within endocrine islets are fitted to be close to 1, i.e., showing almost no correlation consistently over the three time points. Typical results are illustrated in Table 2. One plausible explanation for the observed lack of correlation is that the selected genes are distinctive of different cell types. Within the forming pancreatic islets at this developmental stage, there seem to be no evident clustering effect between distinct cells types bringing them physically close to each other.

**Table 2:** Spatial correlation *γ_ij_* (*mean ± std*) at age 80, 87 and 117 days after fertilization.

| Correlation<br>Intensity |  | Day 80 | Day 87 | Day 117 |
| --- | --- | --- | --- | --- |
| Typical | **SST ↔ INS** | $1.00 \pm 0.02$ | $1.00 \pm 0.01$ | $1.00 \pm 0.01$ |
| | **INS ↔ MUC6** | $1.00 \pm 0.04$ | $1.00 \pm 0.03$ | $0.94 \pm 0.03$ |
| | **INS ↔ ARX** | $1.00 \pm 0.16$ | $1.00 \pm 0.04$ | $0.89 \pm 0.05$ |
| | **ARX ↔ MUC6** | $1.00 \pm 0.03$ | $1.07 \pm 0.03$ | $1.00 \pm 0.04$ |
| Strongest | **EPCAM ↔ PROM1** | $1.26 \pm 0.08$ | $1.26 \pm 0.07$ | $1.33 \pm 0.09$ |
| | **MUC6 ↔ EPCAM** | $1.15 \pm 0.03$ | $1.17 \pm 0.08$ | $1.12 \pm 0.02$ |
| | **MUC6 ↔ PROM1** | $1.09 \pm 0.02$ | $1.13 \pm 0.09$ | $1.19 \pm 0.04$ |

For three pairs of genes a positive spatial correlation was measured at all three time points: EPCAM ↔ PROM1, MUC6 ↔ EPCAM and MUC6 ↔ PROM1. The values are listed in Table 2. As described above, both EPCAM and PROM1 (CD133) are markers of stemness and thus it is not surprising to see their expressions highly correlated spatially, perhaps on the same cell. The positive spatial correlation observed between MUC6 and stem cell markers is certainly interesting because it may indicate a potential role of MUC6 in the differentiation of precursor cells into endocrine cells.

## Discussion

Most in situ transcriptomic studies have so far focused on identification and localization of specific cell types in different organs, mapping data obtained by single-cell RNA sequencing back to tissue sections. Fewer studies have focused at identifying the relations in gene expression between cell types or to other structural and morphological features of the tissue ^12,28,29^. We describe a general analysis tool for spatial correlations of gene expression and carry out temporal study of in situ sequencing data on human fetal pancreas at three developmental ages. We increase the efficiency of the method by probing multiple sites on each transcript and adopting a combinatorial hybridization readout.

A density profile-based method is proposed to study the distribution of transcripts in relation to tissue structures and a statistical model is built to study the spatial correlation between transcripts. The difference between the profiles of each transcript allows us to identify two groups of genes. Notably, we are able to analyze the profiles at different time points and observe how clusters of genes markedly separates from each other. Analyzing samples at three time points, we are able to capture the temporal distribution of single genes within the clusters. We show that MUC6 distribution profile becomes more similar to the group of genes containing endocrine markers and this may indicate a previously unknown role of this gene in the development of pancreatic endocrine cells. The role of mucins genes in the fetal development of several human organs is already known ^30^. Also, MUC6 expression has been identified as an early event in certain pancreatic cancers^31,32^. Our spatial analysis shows that MUC6 distribution positively correlates with other stemness genes and its gene expression clusters with forming endocrine islets following a temporal trend. Altogether these observations identify MUC6 as a candidate marker gene of endocrine differentiation. Notably, other genes of the mucins gene family are present in our panel, but none show strong spatial correlation with endocrine cell or stemness markers. Our density profile-based method is a powerful tool to identify genes of interest at a whole-tissue level.

We show that we can increase the resolution of the spatial analysis by applying our statistical model to genes expressed within clusters of endocrine cells. We find that most gene expressions within identified clusters of endocrine cells are not correlated with each other at the examined time points. Among the pairs of genes with strongest correlations we find epithelial and stemness-related markers EPCAM and PROM1, and MUC6, reinforcing the hypothesis of a role of this gene in cellular differentiation. Because in our analysis we specifically looked for short-range interactions (20 μm) it is possible that these genes are co-expressed or expressed from a niche of progenitor cells. On the contrary, hormones secreting cells identified by expression of GLUC, INS or SST show no correlation with each other at this distance, as expected.

In conclusion, we present a novel method to analyze spatially-resolved transcriptomic dataset which is widely applicable to different technologies and applications. We describe a novel way to explore gene expression data which can be now produced in high throughput by a number of imaged-based techniques. For instance, we demonstrate our method on in situ sequencing data, but the same analysis is applicable to other FISH-based assays. Developmental biology is an ideal application for spatially and temporal- resolved transcriptomic analysis and we demonstrate that our tool can be used to explore and identify potentially novel gene expression patterns and temporal changes.

### Code Availability

We have released a Python implementation of our profile-based method and statistical model analysis on GitHub (https://github.com/RuishanLiu/Gene-Spatial). Most figures in this paper can be reproduced with the codes and datasets in the GitHub repository.

## Acknowledgements

All the imaging was carried out at the Cell Science Imaging Facility, SOE Shriram Center, Stanford University. We thank Cedric Espenel for the discussions on image acquisition and analysis. R.L. is partially supported by Stanford Graduate Fellowship. J.Z. is supported by a Chan-Zuckerberg Investigator grant and by National Science Foundation grant CRII 1657155. M.M. is supported by the Swedish Research Council grant 2015-00599.

## Author Contributions

J.Z., M.M and S.R.Q. supervised the research. J.Z., S.R.Q and M.E. conceived the study. R.L. designed the analysis tool, implemented the algorithm and carried out the computational study. M.M. performed experiments and image analysis. R.J. contributed to the interpretation of the results. S.K.K. procured the samples. R.L., M.M. and J.Z. wrote the manuscript with input from all the authors. All of the authors reviewed the manuscript.

## Competing Interests

The authors declare no competing financial or non-financial Interests.

